# Resolving the structure of phage-bacteria interactions in the context of natural diversity

**DOI:** 10.1101/2021.06.27.449121

**Authors:** Kathryn M. Kauffman, William K. Chang, Julia M. Brown, Fatima A. Hussain, Joy Y. Yang, Martin F. Polz, Libusha Kelly

## Abstract

Microbial communities are shaped by viral predators^1^. Yet, resolving which viruses (phages) and bacteria are interacting is a major challenge in the context of natural levels of microbial diversity^2^. Thus, fundamental features of how phage-bacteria interactions are structured and evolve in “the wild” remain poorly resolved^3, 4^. Here we use large-scale isolation of environmental marine *Vibrio* bacteria and their phages to obtain quantitative estimates of strain-level phage predator loads, and use all-by-all host range assays to discover how phage and host genomic diversity shape interactions. We show that killing in environmental interaction networks is sparse - with phage predator loads low for most bacterial strains and phages host-strain-specific in their killing. Paradoxically, we also find that although overlap in killing is generally rare between phages, recombination is common. Together, these results indicate that the number of hosts that phages infect is often larger than the number that they kill and suggest that recombination during cryptic co-infections is an important mode of phage evolution in microbial communities. In the development of phages for bioengineering and therapeutics it will be important to consider that nucleic acids of introduced phages may spread into local phage populations through recombination, and that the likelihood of transfer is not predictable based on killing host range.

---

Phages are important predators of bacteria – they shape the structure, function, and evolution of natural microbial communities, and they are potential tools to manipulate microbial communities for industrial, bioengineering, and therapeutic applications^5^. Key to understanding the role of phages in natural communities, and to their design and use as efficient and robust tools, is knowledge of their host range in the context of the systems in which they exist or will be used^3^. Yet, how phage host ranges are structured in complex microbial communities remains challenging to address^4^ because the local genomic diversity of phage and bacterial strains is high and, akin to antibody-antigen interactions^6^, phage-bacteria interactions are highly specific. Diverse and powerful approaches have been used to predict phage-bacteria interactions, including: bioinformatic identification of prophages in bacterial genomes^7^; identification of shared sequences or sequence features between phage and bacterial genomes including tRNAs, CRISPR-spacers, and k-mer profiles^8, 9^; single cell sorting and sequencing, which can link host sequences to those of infecting viruses^10^; and viral tagging, in which fluorescently labeled viruses are incubated with hosts and the “tagged” hosts sorted out for sequencing^11–13^. However, no studies since those of Moebus & Nattkemper in the 1980s^14^ have used all-by-all killing cross-tests of cultivated environmental phages and bacterial strains at sufficient scale to resolve a full landscape of potential interactions across multiple levels of phylogenetic diversity, and unfortunately sequencing was not an option at the time of those studies. Analyses of the structure of the Moebus-Nattkemper matrix have provided important insight that there are differences between environmental and laboratory phage-bacteria interaction networks^2, 15^, yet a broad understanding of how genomic diversity shapes interactions between phages and bacteria in nature remains lacking. Here, we uncover how natural levels of phylogenetic diversity shape the structure of interaction networks between phages and bacteria in microbial communities by combining cultivation- and genome-sequencing based approaches to analyze the largest environmental phage-bacteria model system available to date.

To quantify phage predation on closely related bacteria in the environment, we focused on the well-characterized^16–19^ coastal marine heterotrophic Vibrionaceae bacteria as a model system. We isolated >1200 strains, predominantly in the genus *Vibrio*, over three days (ordinal day 222, 261, and 286) during the course of the 3-month 2010 Nahant Collection Time Series^20^ and sequenced the housekeeping gene *hsp60* to initially resolve their phylogenetic relationships. Using 1287 of these isolates as “bait” we quantified concentrations of lytic phages present for each strain in seawater collected on the same days. By using quantitative direct plating agar overlay methods^21, 22^ with virus concentrates^23^, rather than enrichments, we were able to obtain estimates of concentrations of co-occurring lytic phages for each bacterial strain, with 285/1287 (22%) bacterial strains being positive for phage killing. These bait assays provided phage colonies (plaques) from which phage strains could be isolated^24^ to examine their host ranges. Selection of one phage from each plaque-positive host for further study allowed isolation and genome sequencing of a subset of 248 independent phage isolates. Host ranges of each of these phages was assayed against a panel of 294 genome-sequenced bacterial strains, including all plaque-positive hosts and several additional *Vibrio* strains. This yielded a matrix of 72,912 possible killing interactions between genome sequenced phages and bacteria (Supplementary Data File 1).

Our large-scale bait assay revealed that, at the strain level, lytic phage predation pressure on the majority of coastal ocean *Vibrio* is very low (<67 lytic phage L^-1^; limit of detection) compared to total virus-like-particle concentrations (10^10^ VLP L^-1^) common in coastal marine environments^25^ (Fig. 1). As individual strains of the most abundant species in our study typically occur at concentrations of on average <1 cell ml^-1^ ^26^, these findings indicate that encounter rates should be very low between these phages and their *Vibrio* hosts. These observations suggest that mechanisms that increase encounter rates between phages and their hosts - such as host blooms, spatial structure on small scales, and broad host range - should be important features of phage-bacteria interactions in systems where individual host strains are rare; and we therefore explored these features in our matrix.

**Figure 1.**
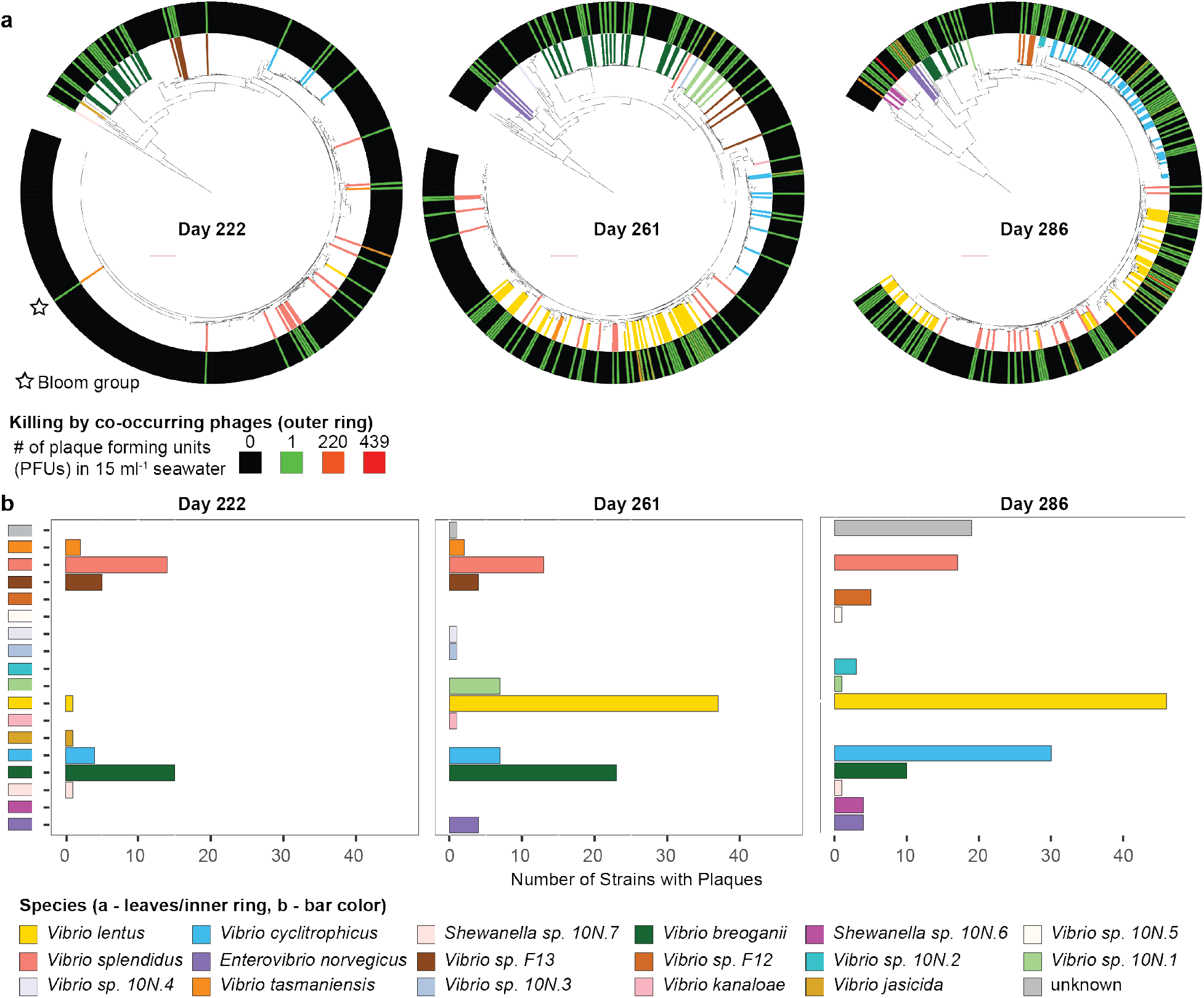
Phage predation pressure on individual bacterial strains appears low overall, is not uniform among closely related bacterial isolates, and varies across days of sampling. **a,** Phylogenetic relationships between Vibrionaceae strains isolated on each of three days and screened for sensitivity to phages in seawater concentrates from the same days (449 bacterial isolates on ordinal day 222; 443 on day 261; 395 on day 286; shown as *hsp60* gene trees with leaves colored by host species for isolates with sequenced genomes). Sensitivity to phage killing is shown in the outer ring, with colors representing the number of plaques formed on each strain. The majority of isolates screened had phage predator loads below the limit of detection (1 plaque forming phage unit (PFU) 15 ml^-1^ seawater); with maximum plaques per strain of 90 PFU ml^-1^ on day 222; 81 on day 261; 430 on day 286. The sole killed representative of an apparent bloom (*Vibrio tasmaniensis* 10N.222.48.A2) is labeled with a star; 28% (125/449) of day 222 isolates were of this *hsp60* genotype and it was not observed on days 261 or 286. Tree scale of 0.1 substitutions per site indicated by red bars. **b,** Strains killed by co-occurring phages (43 plaque-positive on ordinal day 222; 101 on day 261; 141 on day 286) were targeted for genome sequencing and used in subsequent host range assays. Underlying data provided in Supplementary Data File 1 and Source Data Figure 1, see Methods for strain sets analyzed.

The structure of the phage-bacteria interaction network in this study indicates that both broad and narrow host range strategies are important in the coastal marine environment. This emerged when considering the overall structure of the network, where we found that the lytic infections (1436 total) could be organized into 89 discrete groups (modules), each comprised of phages and bacteria interacting largely only with other members of the same module (Fig. 2a, BiMat^27^ modularity measure Q_b = 0.7306). Whereas the three largest modules represented the majority of killing (53% of all interactions, 768/1436 total infections), the majority of modules were “singletons” comprised of only a single phage and bacterial strain interacting exclusively with each other (4% of all interactions, 61/89 modules).

**Figure 2:**
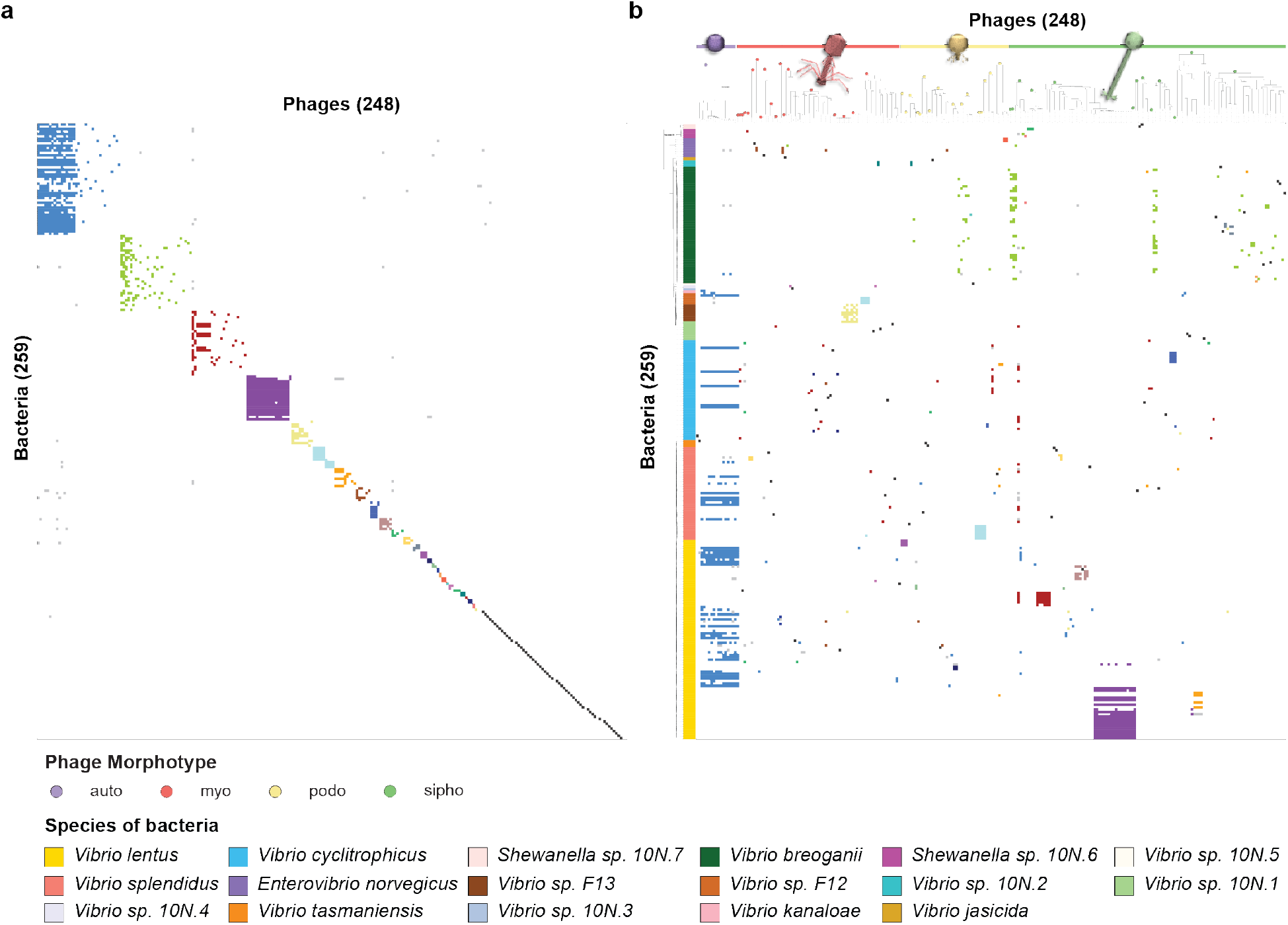
The nested-modular structure of environmental phage-host interaction networks reflects multiple drivers. **a**, Network analysis of the Nahant Collection infection matrix shows an overall nested-modular interaction structure and abundance of one-to-one infections. **b**, Re-organization of the interactions in light of host phylogenetic and phage genomic diversity reveals that modular structure reflects influence of host species, phage genera, phage host range strategies, and bloom dynamics. In both panels bacteria are represented as rows and phages as columns; both panels show the same 248 phages and the subset of 259 Nahant Collection hosts which were infected in the host range assay and for which genomes were also available; interactions in both panels are colored according to BiMat leading eigenvector modules; in panel b bacterial strains are ordered based on phylogeny of concatenated single copy ribosomal protein genes, with leaf colors representing species; in panel b phages are ordered based on manual sorting of VICTOR genus-level trees into groups by morphotype irrespective of their higher order clustering (where genera of different morphotypes can be intermingled, see Methods), each of the 49 genera are represented as a distinct group indicated by a circle filled with the color representing the morphotype of the genus (purple: non-tailed; red: myovirus; yellow: podovirus; green: siphovirus). Underlying data are provided in Supplementary Data File 1, see Methods for strain sets included in the analyses. Phage icon source: ViralZone www.expasy.org/viralzone, Swiss Institute of Bioinformatics.

Central to the organization of each of the three largest modules were phages that were able to kill numerous genomically diverse host strains (Fig. 2b, Extended Data Fig. 1, Supplementary Data File 1). The largest module was organized around killing by members of a new family of recently described phages, the *Autolykiviridae*, whose members can infect some but not all host strains in up to 6 species^28^. The second largest module was likewise organized around phages that killed multiple host strains within a single phylogenetically divergent species, *Vibrio breoganii*. This species is non-motile, lives predominantly attached to macroalgal detritus, and is specialized for degradation of algal polysaccharides^29, 30^ - and thus is also ecologically distinct from other vibrios. The genomically diverse phages infecting *V. breoganii* hosts were nearly all exclusive to this host species in their infections, suggesting that divergence in bacterial ecology is also reflected in interactions with different groups of phages. The third largest module was organized around a single broad host range siphovirus that infected select host strains in 6 species in our network, including members of both the Vibrionaceae and the Shewanellaceae. All three of these large modules, while organized around broad host range phages that could infect multiple specific host strains, included other phages that were effectively entrained into the module as a result of sharing a host strain with the module-defining broad host range phages.

The striking dominance of singleton modules in this network highlights the prevalence and success of exquisitely narrow host range as a strategy in the coastal marine environment. The presence in the network also of modules that contain multiple closely related phages and hosts suggests that blooms may be important in sustaining these otherwise low-encounter singleton interactions. For example, in the 4th largest module (23% of all infections) the majority of phages (18/19) were of >99% pairwise average nucleotide identity (ANI) and infected largely the same set of closely related host strains (18/19 host strains in the module showing >99.95% pairwise ANI). The potential for *Vibrio* to form such blooms is well supported as they proliferate quickly in response to nutrient pulses and have been observed to rapidly undergo large increases in relative abundance in microbial communities in the environment^20, 31^.

We next sought to understand how the observed host range profiles in our matrix were structured in relation to phage morphotypic and genomic diversity. We had previously predicted the morphotypes of these phages using Virfam^24^, and here we used the VICTOR genome classifier tool^32^ to cluster phages and assign them operationally to VICTOR species rank clusters and VICTOR genus rank clusters (hereafter, simply species and genera). Based on the VICTOR classifier, the 248 phages in this study represent 171 species and 49 genera - including 1 genus of non-tailed phages in the *Autolykiviridae*, and 48 genera of tailed phages (myoviruses: 19 genera; podoviruses: 16; siphoviruses:13) (Extended Data Table 1, Extended Data Fig. 2). To ask whether any of these genera included previously described members, we used vConTACT2^33^ to cluster the Nahant Collection phages with >10,000 previously described phages with available genome sequences in NCBI (Supplementary Data File 2). We found the VICTOR and vConTACT2 genus-level clusters to be largely concordant, and identified 17 Nahant Collection genera that include previously described phages, though none in the same species as Nahant phages. The majority of previously described phages in these genera also infected hosts in either the Vibrionaceae or Shewanellaceae, consistent with previous finding that phage genera are largely specific to host families^32^ (see Supplementary Data File 2 for exceptions).

Considering the relationship between phage morphotype and host range breadth in terms of species killed, we found that there were phages of all morphotypes, including the non-tailed autolykiviruses, that killed host strains in only a single species, in two species, in three or more species, and in two genera (Supplementary Data File 3). Three siphoviruses killed hosts in both families of bacteria represented in this study, however, the limited representation of potential non-Vibrionaceae hosts precludes any conclusion about whether this reflects a broader pattern. Interestingly, there were no myoviruses that killed the ecologically distinctive *V. breoganii*, though 19/49 phage genera (71/248 phages) were of this morphotype and this host species was present on all three isolation days (Fig. 1) and well represented in the host range assay (Supplementary Data File 1). Together, these observations indicate that morphotype is not a reliable indicator of the number of host species a phage will infect but may shape access to hosts with different ecological and habitat associations.

We next considered host range profiles in light of phage species and genera and found that overlap in killing is high between phages within species, but low between phages of different species within genera. This feature is evident in visual evaluation of host range profiles (infection matrices in Fig. 3,a,b,c), and is consistent with the striking diversity of putative receptor binding proteins (RBPs) among phages of different species within genera (protein cluster matrices in Fig. 3a,b,c; Supplementary Data File 4). To evaluate these differences quantitatively, we defined a metric of host profile concordance based on Jensen-Shannon distance between host range profiles represented as normalized binary (killed or not-killed) vectors of host strains (see Methods). With this metric, a concordance value of 1 is equivalent to perfect overlap in the host ranges of any two phages, and a value of 0 represents no shared hosts. At the species level we found that concordance values were generally high (Fig. 3d), with phages in 10 species showing perfect overlap in their host range, including 3 species with member phages isolated on different days (this analysis included 105 phages, representing the 28 species with >1 member). By contrast, within genera, concordance in host range among members was generally low (Fig. 3d), even when calculated using a conservative approach that yields higher estimates of concordance when species in a genus contain multiple members (see Methods).

**Figure 3.**
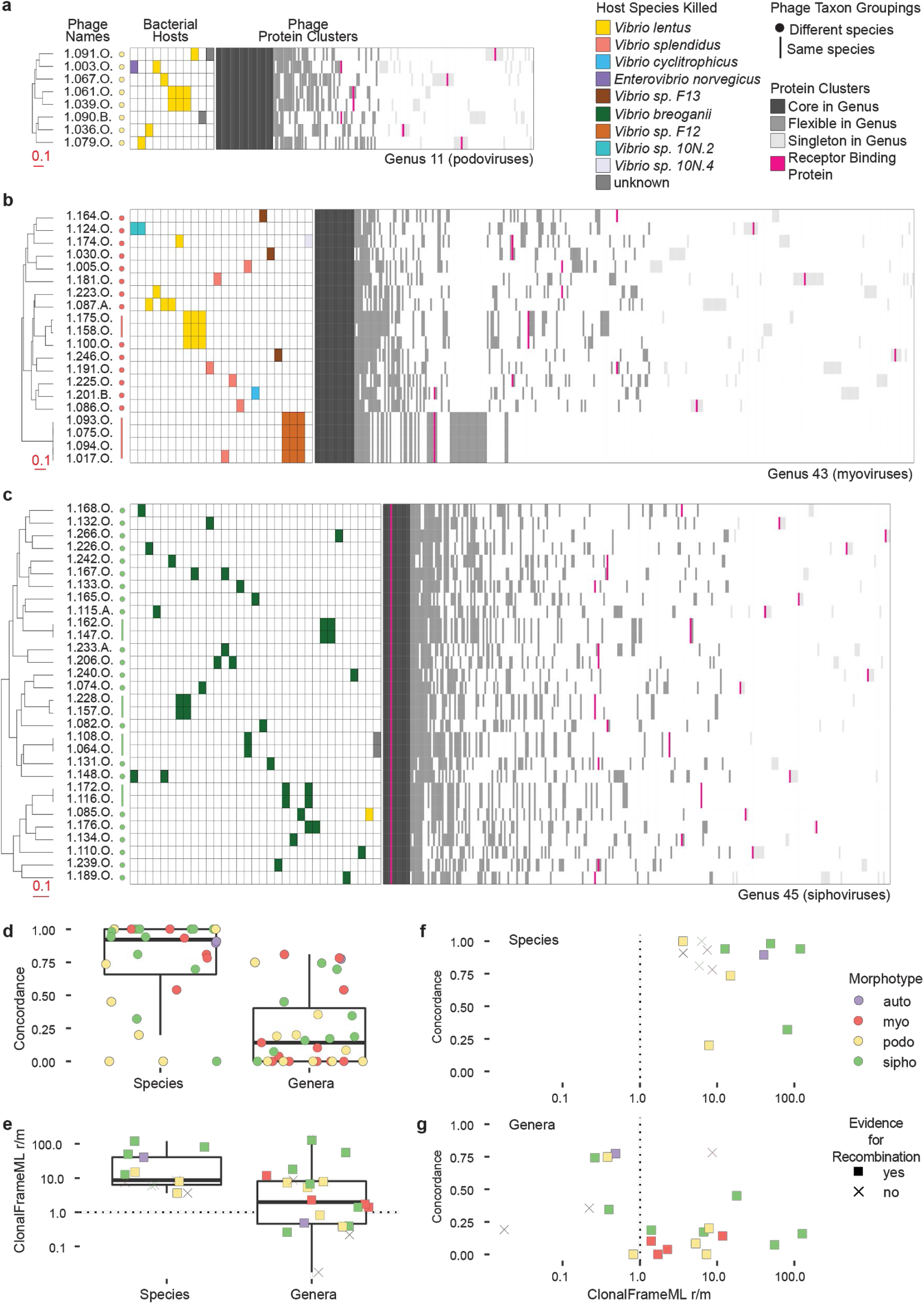
Host range overlap is high within phage species and low within phage genera, but recombination occurs both within and between species within genera. Overlap in host range between phages within genera is low overall but high within species, flexible genes are patchily distributed within genera, and receptor binding proteins are highly diverse; these patterns are observed across morphotypes, as highlighted for genera of: **a**, podoviruses; **b**, myoviruses; and **c**, siphoviruses. Quantified host range profiles for phages across the collection show that: **d**, overlap in killing profiles (concordance) is high within species (28 multi-member species) but low within genera (25 multi-species genera); that **e**, recombination in conserved regions is commonly a greater contributor to genomic diversity in both species and genera; and **f**, that there is no relationship between concordance in killing and recombination for either species or genera. Underlying data and strain information available in Supplementary Data Files 1 and 4, and in Source Data Figure 3, see Methods for description of differences in results when considering only single species representatives in genus-level analyses.

The differences in host range concordance at the phage species and genus levels suggested that there should be corresponding differences in levels of recombination between phage genomes within these groups, yet we found recombination to be occurring both within and between phage species. We observed this qualitatively in the distributions of flexible genes, where in some cases nearest-neighbor phages at the whole-genome level share flexible genes not with each other, but instead with phages of different species within their genus (protein cluster matrices in Fig. 3a,b,c; Supplementary Data File 5). To quantitatively evaluate relative contributions of homologous recombination and mutation (r/m) to diversification we analyzed regions of phage genomes conserved at the species- and genus-levels. At the species level we found that homologous recombination was the greater contributor (r/m >1) to diversification for the majority of species with sufficient members to test (Fig. 3e). This finding of high recombination within phage species is consistent with similar findings in other environments^34, 35^ and indicates that, even in this seemingly rare-encounter opportunity system, phage virion concentrations can reach high enough local concentrations to result in co-infections between members of the same species in cells of their shared hosts. Surprisingly however, we also found a strong signal of recombination (r/m >1) at the genus level (Fig. 3e), even when calculated using a conservative approach that underestimates r/m in genera containing species with multiple members (see Methods).

The importance of recombination at both the species and genus levels indicates that overlap in killing between phages is not predictive of their potential for recombination in the context of natural microbial communities. This is corroborated when both quantitative metrics are considered together, which shows a lack of a positive association between host range concordance and r/m on both the genus (Fig. 3f, Spearman correlation -0.2695786) and species levels (Fig. 3g, Spearman correlation -0.2417582). Altogether, these observations suggest that phages are likely infecting many more hosts than they are able to kill and that therefore co-infections of host cells by multiple different phages (necessary to allow for recombination between phage genomes) are more common than predicted by killing host ranges.

To understand the potential maximal extent of recent gene flow among all phages in this collection we used a k-mer based approach, and found evidence of sequence sharing between phages of different morphotypes with non-overlapping host killing. We first used the liberal metric of occurrence of sharing of any 100% identity 25-base pair (25-mer) length sequences between phages, such 25-mers are sufficiently unique in bacterial genomes to be used to recapitulate strain and species level relatedness^36^, and provide a marker for potential recent horizontal gene transfer events^37^. Using this approach we found far greater potential connectivity of gene flow than suggested by overlaps in host killing (Fig. 4a,b). Considering next the more conservative metric of total numbers of 25-mers shared between any two phage genomes we also found evidence for sequence sharing between divergent phages with non-overlapping host ranges; with the maximum number of shared 25-mers between any pair of phages of different morphotypes being 6,169 (Fig. 4c). Finally, an approach designed to detect specific recent gene transfer events from conserved regions of phage genomes revealed connections between phages in different genera and without overlapping host ranges, and between which there were often long regions of high genomic sequence similarity (Fig. 4d, Extended Data Fig. 3).

**Figure 4.**
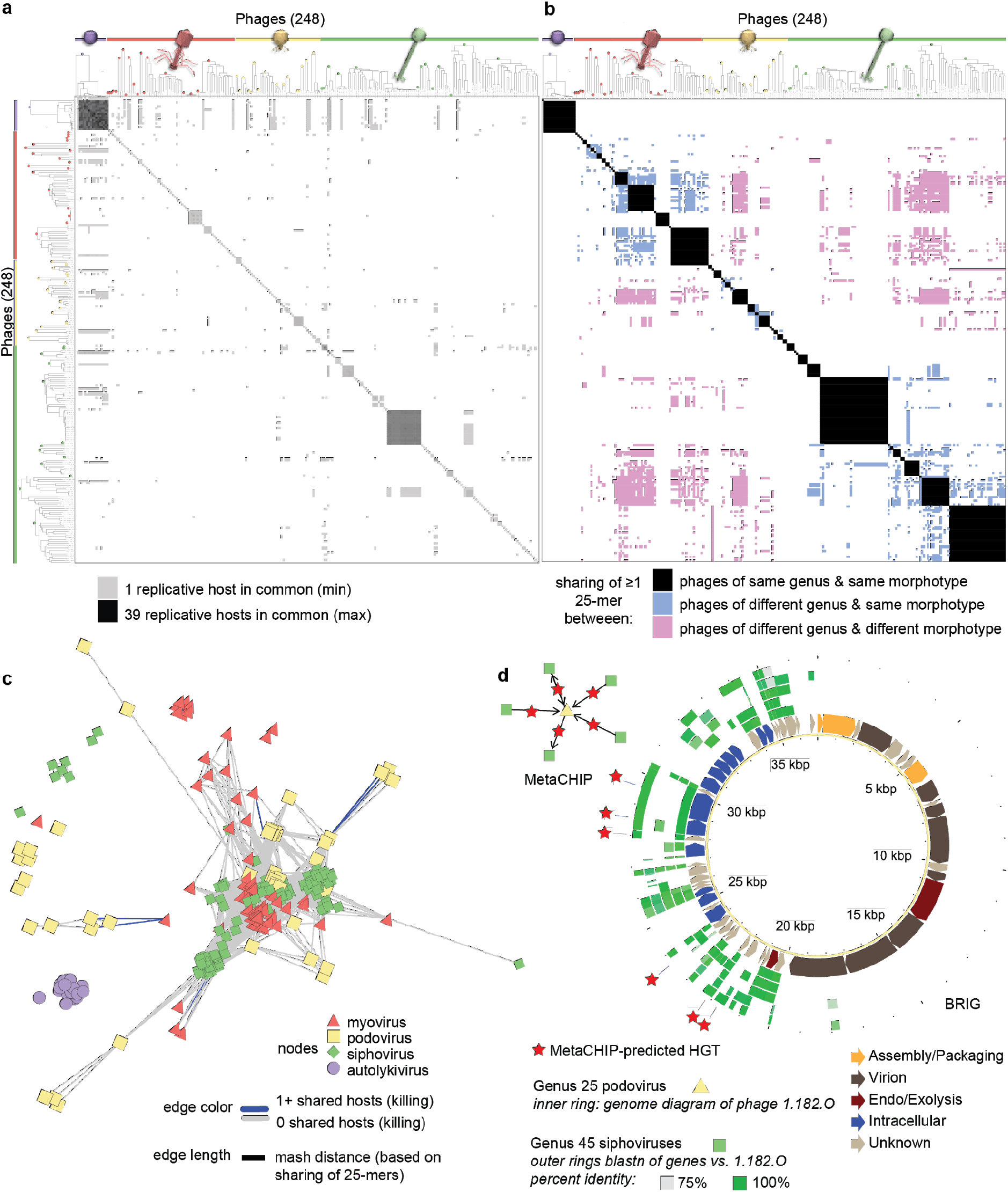
Overlap in killing of host strains is sparse and does not reflect sequence sharing between phages of different genera or morphotypes. Matrix representations of pairwise comparisons between phages, show that: **a**, occurrence of shared replicative hosts, does not reflect **b**, occurrence of sharing of ≥1 25-mer between phages of different genera and morphotypes; in both panels phages are ordered based on manual sorting of VICTOR genus-level trees into groups by morphotype irrespective of their higher order clustering (where genera of different morphotypes can be intermingled, see Methods). Extent of sequence sharing that can occur between phage morphotypes and genera that are largely non-overlapping in host killing is revealed by: **c**, network visualization of phage genome connectivity by Mash distance, which is defined by sharing of 25-mers; and **d**, occurrence of extensive sequence similarity between phage of different morphotypes, as visualized here using BRIG, for which specific directional horizontal gene transfer events can also be detected using MetaCHIP. Underlying data are provided in Source Data Figure 4, see Methods for strain sets included in the analyses. Phage icon source: ViralZone www.expasy.org/viralzone, Swiss Institute of Bioinformatics.

Considering the potential for widespread sequence sharing to influence mapping of viral reads to reference genomes we investigated cross-mapping using a recently developed and powerful k-mer based pseudo-alignment approach. We found that cross-mapping of reads can occur between phages of different species, genera, and morphotypes within this collection. False positive classifications of reference presence were reduced when using shorter simulated read lengths as a result of overall lower collateral sequence coverage when the basis for the mapping was a single k-mer match in the sequence (Extended Data Fig. 4, and see Methods). This observed potential for cross-mapping calls for a cautious approach in using read-mapping to reference genomes in determining whether specific phages are present in metagenomic samples or predicting which hosts virus pools are interacting with.

The overall prevalence of sequence sharing between phage genomes suggests that they harbor homologous recombination systems, and indeed we find that recombinase genes are encoded by the majority of phages in this collection. Low fidelity single strand annealing protein (SSAP) based recombinase systems such as those in the Rad52-superfamily (e.g. *λ* Red, ERF, and Sak) are common in temperate phages^38^, and are thought to play an important role in their extensive genome modularity and mosaicism^39^. Such recombinases have been shown to be associated with large-scale recombination events of up to 79% genome length between incoming temperate phages and resident prophages^40^, and *in vitro* use of phage-derived recombinase systems for genetic engineering (“recombineering”) is a powerful tool for building genetic constructs because they can facilitate recombination between sequence regions with as little as 23-bp homology^41^. Noting a number of putative SSAP recombinase genes in our initial annotations, we sought to more systematically evaluate their representation in our collection of predominantly lytic phages. Using representative sequences^38^ as seeds for iterative searches, followed by gene neighborhood analysis, we identified cryptic putative SSAP-like recombinases in 224/248 (90%) phages, with 117 of these resembling low-fidelity Rad52-superfamily and Sak4-like Rad51-superfamily recombinases commonly associated almost exclusively with temperate phages^38^ (Supplementary Data File 4). These results suggest that just as horizontal gene transfer in microbial communities may allow bacteria to evolve resistance to phages, recombination and genetic exchange between lytic phages may likewise be important in overcoming this resistance.

Altogether, the findings of this work appear at first contradictory – that in natural microbial communities killing is exceedingly sparse but also that recombination between phages is common. Yet, these observations are well reconciled when considered under a unifying conceptual framework that integrates our observations with the emerging understanding of the prevalence and diversity of horizontally acquired mobile phage defense islands^42–44^ that act intracellularly in bacteria to limit killing once phages have injected their genomes. Here, we show that co-occurring phages in the same phage genus are highly diverse in their receptor binding proteins, that recombinase systems are strikingly prevalent in phage genomes, and that there is significant discord between killing host range and observed recombination among phages. Under a unified model, we suggest that diverse phages enter hosts using different receptors, that some of these entering phages are targeted by defense systems that degrade their genomes and thus limit their ability to kill, and that in co-infections with other phages these partially-degraded genomes are available for recombination and yield recombinant phage progeny (Fig. 5).

**Figure 5.**
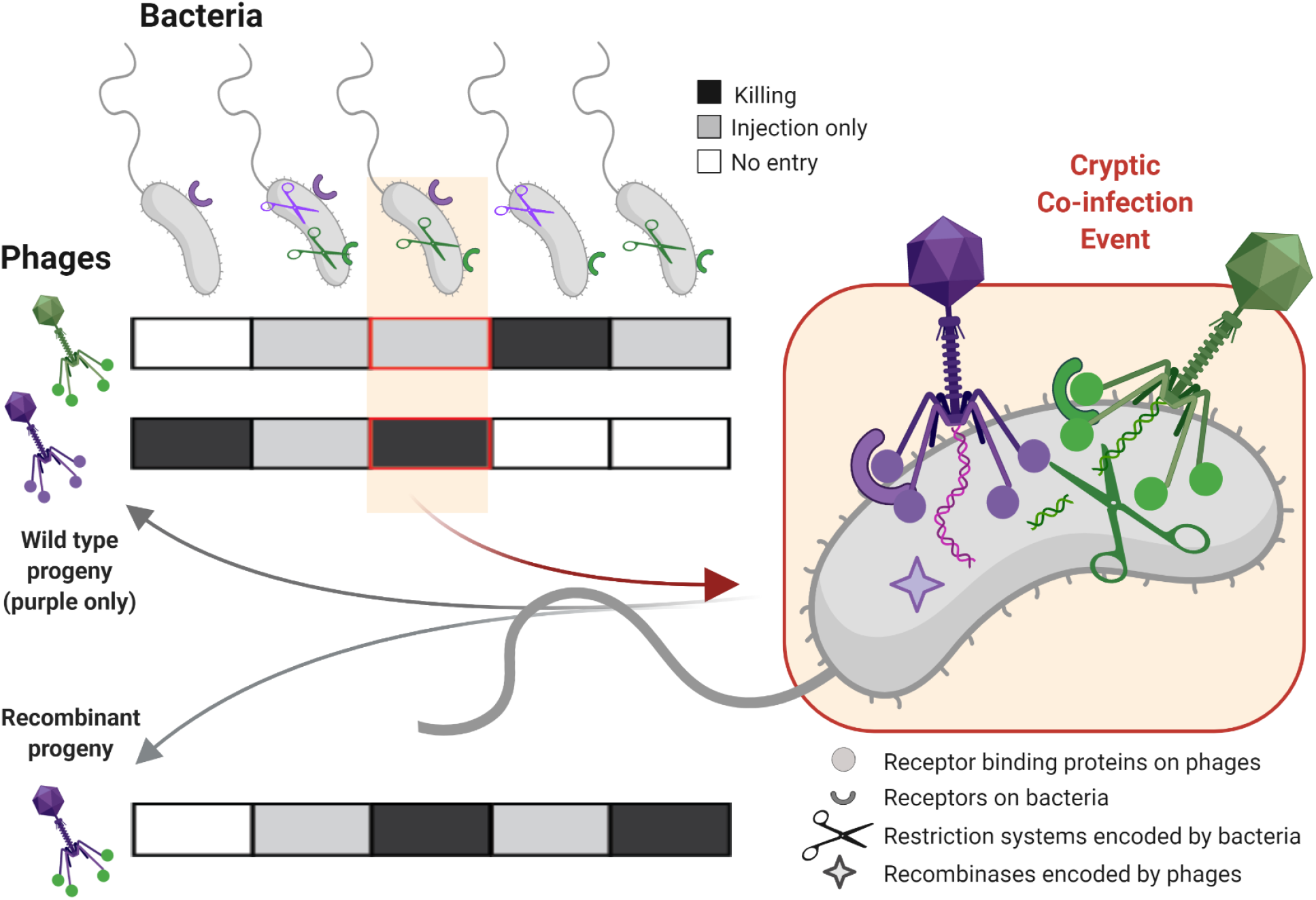
Common cryptic coinfection as a unifying conceptual model for phage-host interactions. There exists a paradox that whereas phages are thought to be narrow host range, their genomes show evidence for high recombinogenicity - this has raised the question of whether both elements are in fact true “in the wild”, and if so, how they can be reconciled. Integrating our work with current understanding we propose the following model to resolve this question: Diverse phages are co-infecting host cells using different receptors and receptor binding proteins, some of these phages are prevented from replicating by intracellular host defenses (and thus appear narrow in host range), but recombination between co-infecting phages (facilitated by recombinases common in phage genomes) allows fragments of thwarted phage genomes to live on in recombinant progeny co-occurring in the environment with their parent strains. Figures created with BioRender.com.

This work highlights the importance of considering that assays of killing host ranges are not predictive of the potential for diffusion of exogenous phage genome constructs into local phage genome pools when they are introduced for bioengineering and therapeutic applications. Infective host ranges of phages are likely far broader than indicated by killing assays, allowing for recombination between co-infecting phages to act as a prevalent mode of lytic phage genome evolution in microbial communities. This importance of cryptic co-infections indicated by our results integrates well with the emerging understanding that complex phage interactions with sub-optimal hosts^43, 45^, and other phages^46–48^, are important forces shaping the dynamics and evolution of phage-host interactions in complex microbial communities.

## Supporting information

Supplementary_Data_File_1.Bacteria_Phage_Infections_Infos

Supplementary_Data_file_2.vCONTACT2_analysis

Supplementary_Data_File_3.Phage_HostTaxa_Summary_Data

Supplementary_Data_file_4.Protein_Annotations

Source_Data_Figure_3.KillingConcordance_and_ClonalFrameML

Source_Data_Figure_4.HostSharing_Mash_MetaCHIP

Source_Data_Extended_Data_Figure_5.FastViromeExplorer

## Acknowledgements

We are grateful to Jan Meier-Kolthoff and Markus Göker, for running their VICTOR tool analysis on the Nahant Collection phage genomes on their cluster. We thank Michael Cutler, Philip Arevalo, and Joseph Elsherbini for support in sequencing, assembly, and annotation of bacterial genomes. We thank David VanInsberghe for support in early phage genome analyses. We thank all members of the 2010 Polz lab for support with field sampling and bacterial isolation, and in particular Michael Cutler, Alison Takemura, Hong Xue, Tara Soni, Gitta Szabo. We thank the Edward Ruby lab for sharing *Vibrio fischeri* strains for inclusion in the host range assay.

## Author contributions

KK and WC designed the study, performed analyses, and wrote the manuscript with substantial contributions throughout from JB, FH, JY, MP, and LK.

## Author information

The authors declare no competing financial interests.

## Code availability

All custom codes associated with this work are available from the authors upon request.

## Data availability

Information about bacteria and phage isolates, infections, and BiMat module assignments are provided in Supplementary Data File 1. Results of vConTACT2 analyses are available in Supplementary Data File 2. Summary data for all operational phage taxa identified in this work are provided in Supplementary Data File 3. Annotations of all proteins and protein clusters of phages in this work are provided in Supplementary Data File 4. Source data underlying Figures 3 and 4, and Extended Data Figure 5, are all also provided as Source Data files. New bacterial genomes and *hsp60* sequence data deposited with this work are included under the Nahant Collection of NCBI BioProject with accession number PRJNA328102.

## Biological materials availability

Phage and bacterial isolates are available from the authors upon request.

**Extended Data Figure 1.**
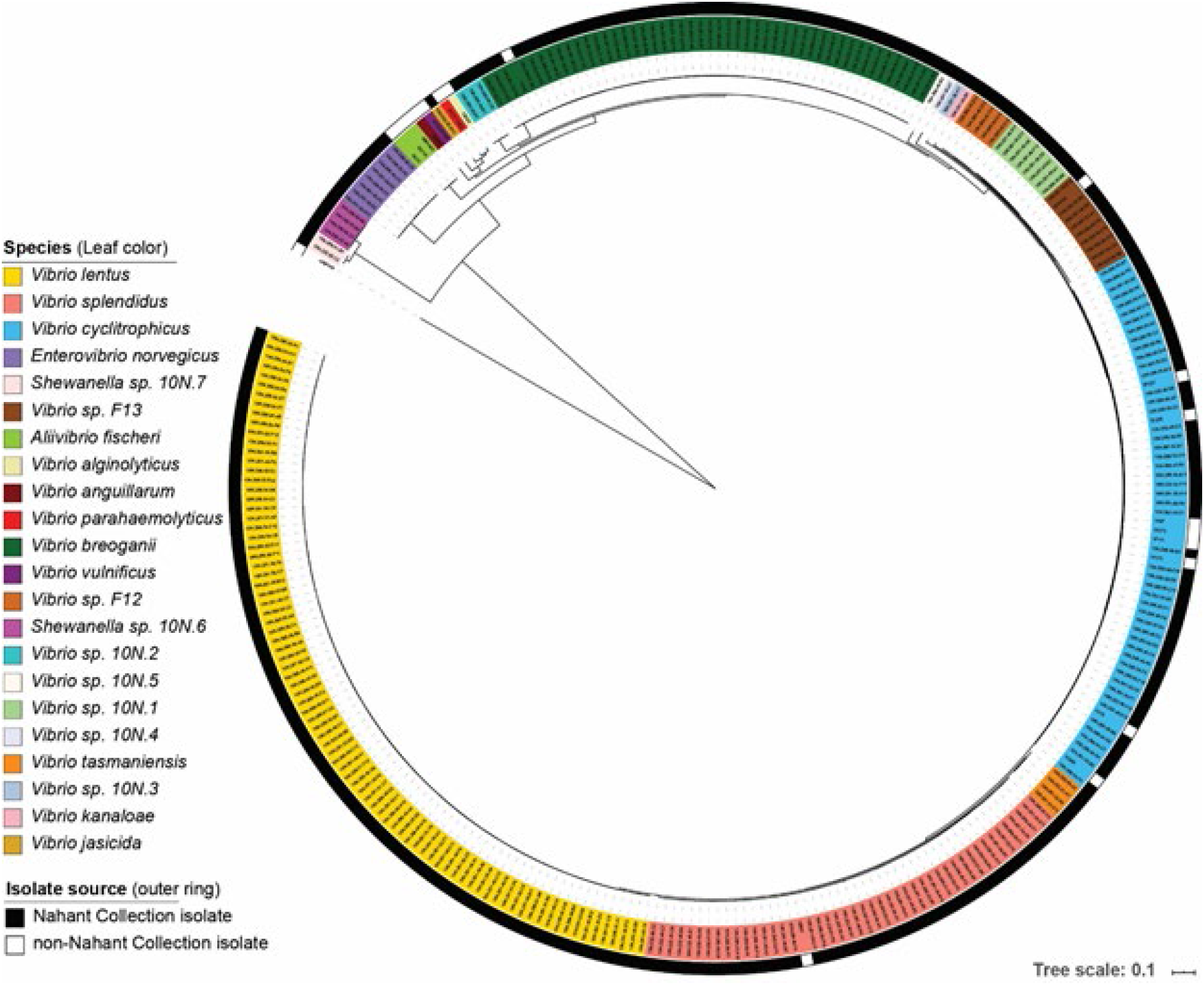
Bacterial isolates included in the large scale host range assay include >45 closely related members of each of 4 *Vibrio* species, as well as representatives of at least 18 other species in the family Vibrionaceae. Tree based on concatenated single copy ribosomal protein genes. Underlying bacterial strain information provided in Supplementary Data File 1.

**Extended Data Figure 2.**
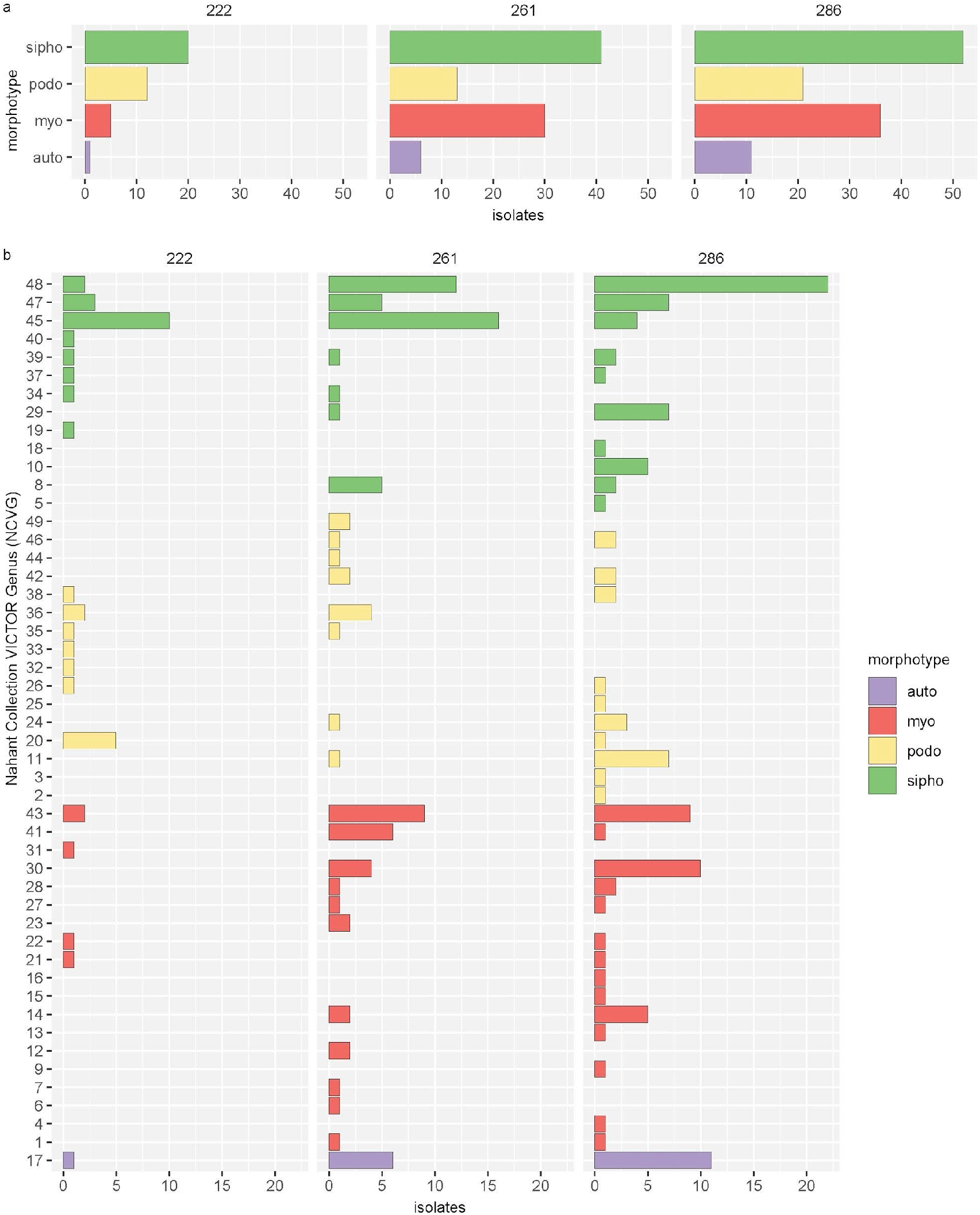
Morphotypes and genera of phages isolated on each of the three sampling days. Underlying phage strain information available in Supplementary Data File 1.

**Extended Data Figure 3.**
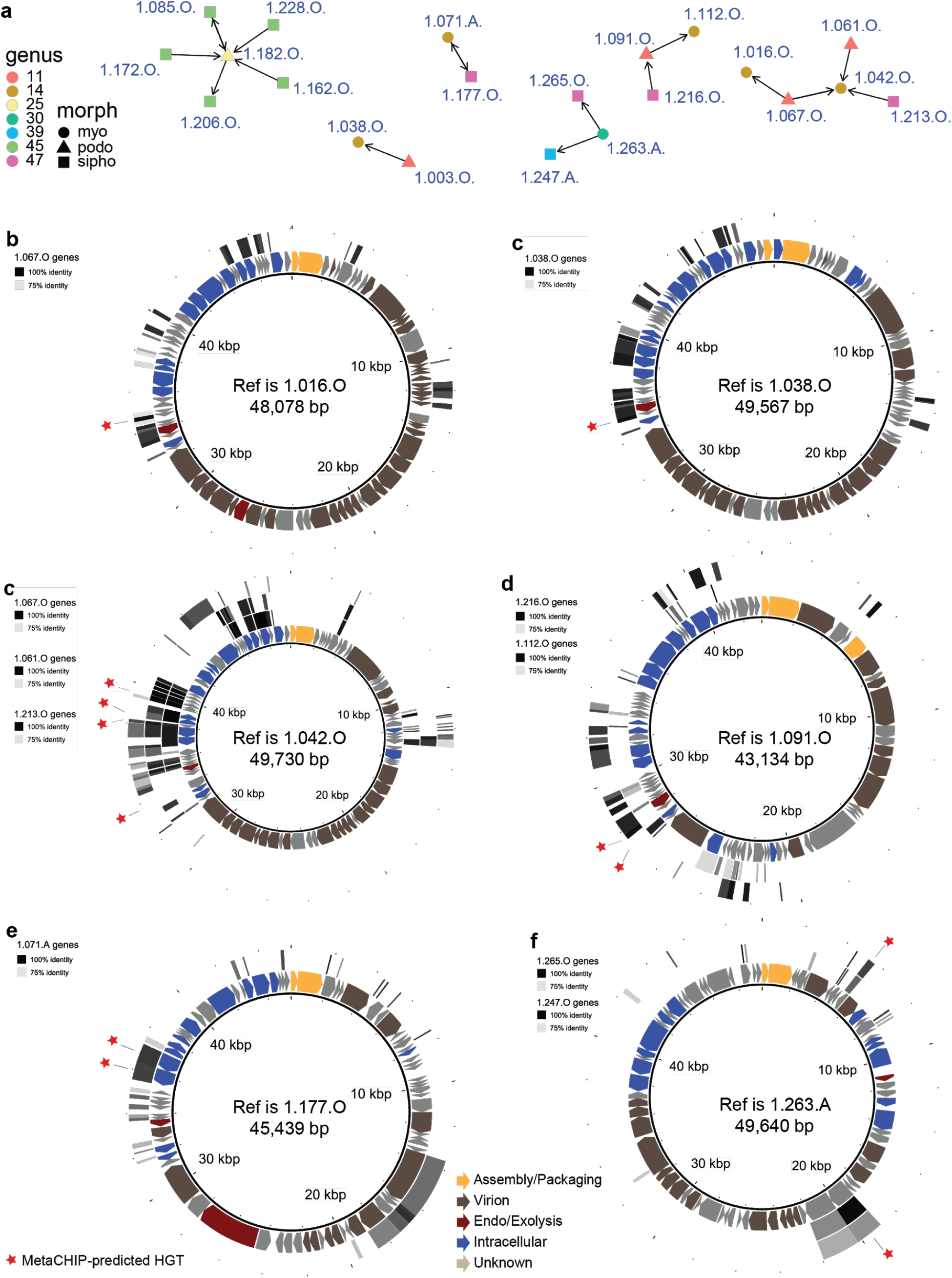
Horizontal gene transfer between phages in different genera and of different morphotypes and with no shared hosts killed. **a**, Candidate gene transfer events between phages in different genera and with different morphotypes were identified by MetaCHIP, which considers conservation with donor groups to predict transfers into recipients. b, Extent of sequence sharing between pairs identified using MetaCHIP was further examined using BLAST-based mapping using BRIG, revealing that there were commonly additional regions of high sequence identity in neighboring genomic regions. Predicted open reading frames in phage genomes are depicted as arrows and colored according to broad gene functional categories; stars indicate sequence sites involved in transfer events predicted by MetaCHIP. Underlying data is provided in Source Data Figure 4 and Supplementary Data File 4.

**Extended Data Figure 4.**
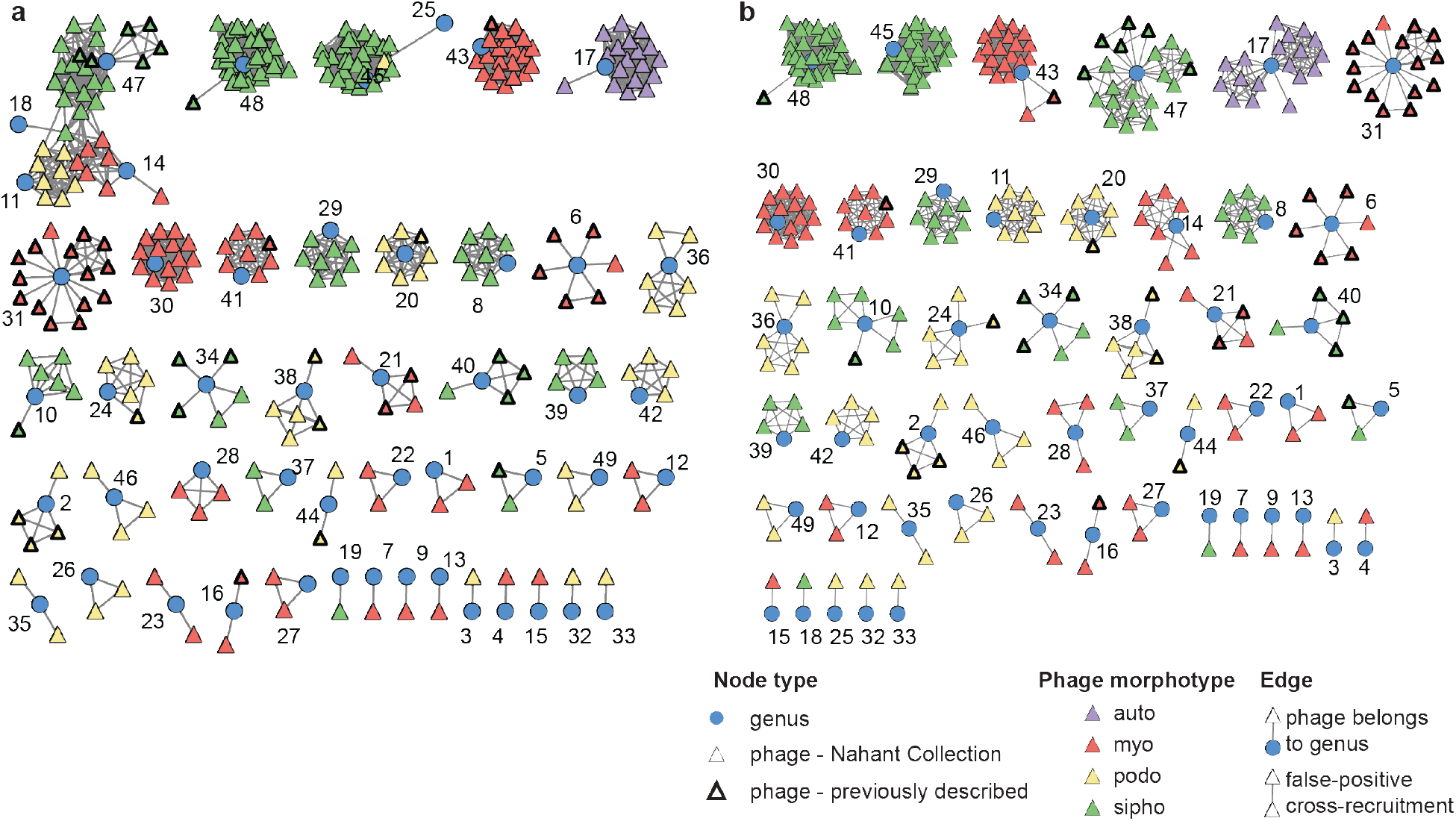
Non-specific recruitment of reads can lead to false positives in mapping of viral reads to reference genomes. Pseudoalignment-based mapping of reads to reference genomes leads to extensive cross-mapping within genera as well as between phages of different morphotypes. Cross-mapping is defined operationally as determination of a virus as “present” by the pseudoalignment-based viral metagenome characterization tool FastViromeExplorer when using default settings and 100,000 simulated reads of either a, 250bp or b, 100bp. Simulated reads from each virus were tested for mapping against all other phages in the dataset individually, including 248 Nahant Collection phages and 47 previously isolated phages identified as members of Nahant Collection genera. As a result of the requirement for mapping of only a single 31-mer to assign a read as mapped, false-positives are more frequently observed when using longer reads as a result of higher overall coverage. Patterns of cross-mapping vary across genera and range from observations of no within-genus cross-mapping to cases of extensive between-genus cross-mapping, including between genera with phages of different morphotypes. Underlying data, including proportion of reads mapped in comparison with mapping to self, are provided in Source Data Extended Data Figure 4.

**Extended Data Table 1.**
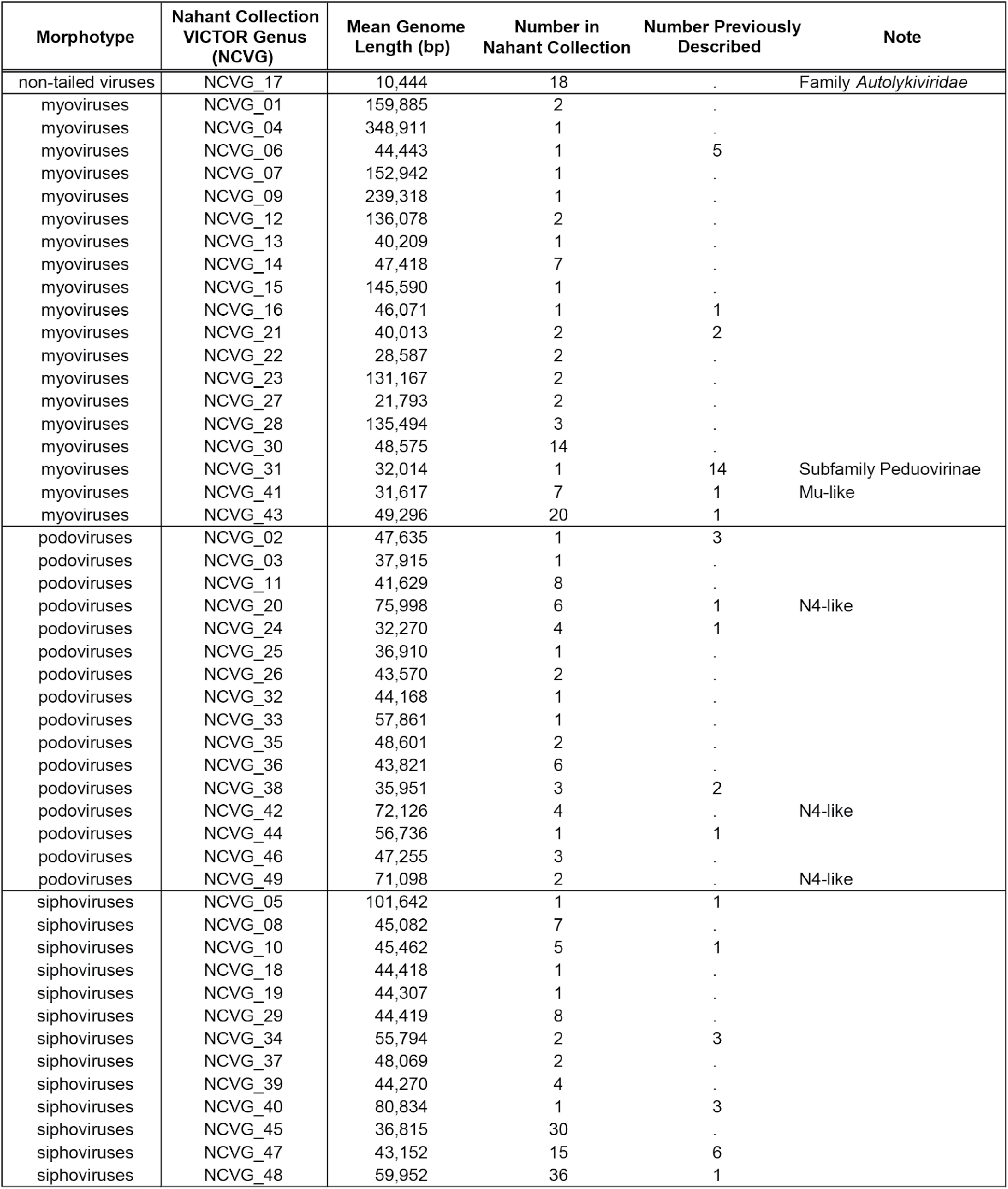
**Overview of Nahant Collection phages by VICTOR genus.**

## Methods

### Sampling

#### Environmental sampling

Samples were collected from the littoral marine zone at Canoe Cove, Nahant, Massachusetts, USA, on 22 August (ordinal day 222), 18 September (261) and 13 October (286) 2010, during the course of the three month Nahant Collection Time Series sampling^20^.

### Bacterial isolation and characterization

#### Bacterial isolation

Bacterial strains were isolated from water samples using a fractionation-based approach^16^ as previously described^24, 28^. In brief, seawater was passed sequentially through 63um, 5um, 1um, and 0.2um polycarbonate filters; material recovered on the filters was resuspended by shaking for 20 minutes; dilution series of resuspended cells were filtered onto 0.2um polyethersulfone filters in a carrier solution of artificial seawater (40g Sigma Sea Salts, S9883; 0.2um filtered), and filters placed directly onto *Vibrio*-selective MTCBS plates (Difco TCBS Agar supplemented with with 10g NaCl per liter to 2% final w/v). Colonies (96) from each of three replicates of each size fraction were selected from the dilution plates with the fewest numbers of colonies (1,152 isolates per isolation day). Colonies were purified by serial passage, first onto TSB-II (Tryptic Soy Broth, 1.5% Difco Bacto Agar, amended with 15g NaCl to 2% w/v), second onto MTCBS, finally onto TSB-II again. Colonies were inoculated into 1ml of 2216 Marine Broth (Difco) in 96-well 2ml culture blocks and allowed to grow, shaking at room temperature, for 48 hours. Glycerol stocks were prepared by combining 100 ul of culture with 100 ul of 50% glycerol in 96-well microtiter plates and sealed with adhesive aluminum foil for preservation at -80°C.

#### Bacterial hsp60 gene sequencing

To obtain *hsp60* gene sequences for isolates, Lyse-N-Go (LNG) treatments of subsamples of the same overnights cultures used in the bait assay (described below) were used directly as template in PCR amplification reactions. PCR reactions were prepared in 30 ul volumes, as follows: 1 ul LNG template, 3 ul 10x buffer, 3 ul 2mM dNTPs, 3 ul 2um hsp60-F primer, 3 ul 2um hsp60-R primer, 0.3 ul NEB Taq, 16.7 ul PCR-grade HOH; with hsp60-F (H279) primer sequence: 5’-GAA TTC GAI III GCI GGI GAY GGI ACI ACI AC-3’, and hsp60-R (H280) primer sequence: 5’-CGC GGG ATC CYK IYK ITC ICC RAA ICC IGG IGC YTT-3’ ^49^. PCR thermocycling conditions were as follows: initial denaturation at 94°C for 2 min; 35 cycles of 94°C for 1 min, 37C for 1 min, 72C for 1 min; final annealing at 72C for 6 min; hold at 10C. PCR products were cleaned up by isopropyl alcohol (IPA) precipitation, as follows: addition of 100 ul 75% IPA to 30 ul PCR reaction product, gentle inversion mixing followed by 25 min incubation at RT, 30 min centrifugation at 2800rcf, addition of 50 ul 70% IPA with gentle inversion wash, centrifugation at 2000rcf, inversion on paper towels to remove IPA, 10 min centrifugation at 700rcf, air drying in PCR hood for 30 min, resuspension in 30 ul PCR HOH. PCR products were Sanger sequenced (Genewiz, Inc.) using hsp60R primer, as follows: 5 ul of 5um hsp60R primer, 7 ul nuclease free water, 3 ul DNA template. For a subset of strains *hsp60* sequences were obtained from subsequently determined whole-genome sequences. *Hsp60* sequences were aligned to the *hsp60* sequence previously published for Vibrio 1S_84 and trimmed to 422 bases using Geneious. Accession numbers for these 1287 strains are provided in Supplementary Data File 1, where they are identified as “baxSet1287”.

#### Bacterial hsp60 phylogenies

A phylogenetic tree of relationships among bacterial isolates screened in the bait assay (described below) was produced based on a 422 bp fragment of the *Hsp60* gene, derived either from Sanger or whole genome sequences; with *E. coli* K12 serving as the outgroup. Sequences from each of the three days of isolation were aligned using muscle v.3.8.31 ^50^ with default settings (*muscle -in $seqsALL -out $seqsALL.muscleAln*), and a single tree including all 1287 sequences from all the days was generated using FastTree v.2.1.8 ^51^ (*FastTree -gtr -gamma -nt -spr 4 -slow < $seqsALL.muscleAln > $seqsALL.muscleAln.fasttree*). For presentation in Fig. 1 three sub-trees including only nodes from each day were produced using PareTree v.1.0.2 ^52^ (*java -jar PareTree1.0.2.jar -t O -del notDay222.txt -f$seqsALL.$round.muscleAln.fasttree.DAY222*). Trees were visualized using iTOL^53^ and painted with metadata for each of the strains, including: sensitivity to killing by co-occurring lytic phage predators collected on the same day and, for the subset of strains that were genome sequenced and also included in the host range matrix, the bacterial species, based on concatenated ribosomal protein analysis using RiboTree ^54^ as described below. Isolation days for each of the strains included in these analyses are provided in Supplementary Data File 1, where these strains are identified as “baxSet1287”.

#### Bacterial genome sequencing and assignment to populations

To assign genome-sequenced bacterial isolates used in the host range assay to species, we use the RiboTree tool ^54^ to produce a phylogeny based on concatenated single copy ribosomal proteins as in^29^. We include strains of previously described Vibrionaceae in preliminary analyses as reference strains and assign species names to new isolates based on clustering with named representatives, as well as provide placeholder names for newly identified clades with no previously described representatives. Trees were visualized using iTOL^53^ and the representation including only those strains included in the host range assay is shown in Extended Data Figure 1; population assignments and accession numbers for this set of 294 genomes, which also includes a small number of previously isolated bacterial strains that were included in the host range assay (described below), are provided in Supplementary Data File 1, where they are identified as “baxSet294”.

### Viral isolation and characterization

#### Viral sample collection

The iron chloride flocculation approach was used to generate 1000-fold concentrated viral samples from 0.2um-filtered seawater, as follows. For each isolation day, triplicate 4L seawater samples were filtered through 0.2 Sterivex filters into collection bottles, spiked with 400uL of FeCl3 solution (10gl^−1^ Fe; as 4.83g FeCl_3_•6H_2_O into 100ml H_2_O), and allowed to incubate at room temperature for at least 1 hour. Virus-containing flocs were then recovered from the sample by filtration onto 90mm 0.2um polycarbonate filters (Isopore, GTTP09030, Millipore) under gentle vacuum; filters were folded into quarters under vacuum and inserted into a 7ml borosilicate glass vial. A volume of 4ml of oxalate-EDTA solution (prepared from stock solution as 10ml 2M Mg_2_EDTA, 10ml 2.5M Tris-HCl, 25ml 1M oxalic acid; adjusted to pH with 10M NaOH; final volume 100ml; used within 7 days of preparation and maintained at room temperature in the dark) was added to the vial and the sample allowed to dissolve at room temperature for at least 30 minutes before transfer to storage at 4°C. A reagents used in this original formulation (JT-Baker 7501 Mg_2_EDTA) is no longer available and an updated recipe is provided elsewhere^55^.

#### Bait assay and associated viral plaque archival

In order to obtain quantitative estimates of co-occurring phage predator loads at bacterial strain level resolution, and generate plaques from which to isolate phages, we exposed approximately 1440 purified bacterial isolates to phage concentrates from their same day of isolation as follows. Bacterial strains screened included 480 isolates from each ordinal day, representing 120 strains from each of 4 size-fractionation classes (0.2 um, 1.0 um, 5.0 um, 63 um) details of isolation origin are provided for each strain in Supplementary Data File 1, and description of naming conventions is as previously described^24^. For the bait assay each strain was mixed in agar overlay with seawater concentrates containing viruses (15 ul concentrate, equivalent to 15 ml unconcentrated seawater; derived from pooling of three replicate virus concentrates from each day). Agar overlay were performed using the Tube-free methods as previously described^22^, with the following modifications. Bacterial strains were prepared for agar overlay plating by streaking out from glycerol stocks onto 2216MB agar plates with 1.5% Bacto Agar (Difco), and allowed to grow for 2 days at room temperature.

Strains were then inoculated into 1 ml 2216MB in a 96-well culture block and incubated 24 hours at room temperature shaking at 275rpm on a VWR DS500E orbital shaker. Immediately prior to use in direct plating the OD600 was measured in 96-well microtiter plates and subsamples were taken for Lyse-N-Go processing for DNA (10 ul culture, 10 ul LNG). Phage concentrates were prepared for plating by pooling 1.2ml from each of the concentrate replicates into a 7ml borosilicate scintillation vial. Cultures were transferred from overnight culture blocks to 96-well PCR plates in 100 ul volume and 15 ul of pooled phage concentrate was added to cultures one row at a time, with each row plated in agar overlay before adding phage concentrate to the next row of bacterial cultures. Mixed samples of 10 ul bacterial overnight culture and 15 ul pooled phage concentrate were transferred to the surface of bottom agar plates (2216MB, 1% Bacto Agar, 5% glycerol, 125 ml l^-1^ of chitin supplement [40 g L^-1^ coarsely ground chitin, autoclaved, 0.2 um filtered]). A 2.5 ml volume of -52°C molten top agar (2216MB, 0.4% Bacto Agar, 5% glycerol) was added to the surface of the bottom agar and swirled around to incorporate and evenly disperse the mixed bacterial and phage sample into an agar overlay lawn. Agar overlay lawns were held at room temperature for 14-16 days and observed for plaque formation. Glycerol was incorporated into this assay to facilitate detection of plaques (Santos et al., 2009). Chitin supplement was incorporated into this assay to facilitate detection of phages interacting with receptors upregulated in response to chitin degradation products. A variety of preliminary tests exploring potential optimizations to agar compositions for direct plating indicated that the addition of chitin did not negatively impact recovery of plaques with control phage strains tested. After approximately 2 weeks, plaques on agar overlay lawns were cataloged and described with respect to plaque morphology and plaques were picked for storage using the Archiving Plaques method as previously described^22^. All plaques were archived from plates containing less than 25 plaques, on plates with larger numbers of plaques a random subsample of plaques from each distinct morphology were archived. A polypropylene 96-well PCR plate was filled with 200 ul aliquots of 0.2 um filtered 2216MB, agar plugs were collected from plates using a lml barrier pipette tip and ejected into the 2216MB, skipping one well between each sample to minimize potential for cross-contamination, for a final count of 48 phage plugs per plate. Plaque plugs were soaked at 4°C for several hours to allow elution of phage particles into the media. After soaking, 96-well plates were centrifuged at 2,000 rcf for 3 minutes before proceeding to the next step. Plug soaks were then processed for two independent storage treatments. For storage at 4°C, plates were processed by transferring 150 ul of eluate from each well to a 0.2um filtration plate (Millipore, Multiscreen HTS GV 0.22um Filter Plate MSGVS22) and gently filtered under vacuum to remove bacteria, the cell-free filtrates containing eluted phage particles from each plaque plug were stored at 4°C. For storage at -20°C, 50 ul of 50% glycerol was added to the residual ∼50 ul of the plug elution, often still containing the agar plug. In this way all plaques were characterized and many plaques from each strain were archived in two independent sets of conditions. Total plaque counts for all strains included in the bait assay are represented in Figure 1, and provided in Supplementary Data File 1, where they are identified as “baxSet1287”.

#### Viral purification

A subset of plaques archived during the bait assay was selected for phage purification, genome sequencing, and host range characterization. This subset included single randomly-selected representatives from each plaque-positive bacterial strain. Minor details of the purification and lysate preparation varied across samples but were largely as follows. Phages were purified from inocula derived primarily from -20°C plaque archives, and secondarily from 4°C archives when primary attempts with -20°C stocks failed to produce plaques. Three serial passages were performed using Molten Streaking for Singles^22^ method. Agar overlay lawns for passages were prepared by aliquoting 100 ul of host overnight culture (4ml 2216MB, colony inoculum from streak on 2216MB with 1.5% Bacto Agar, shaken overnight at RT at 250rpm on VWR DS500E orbital shaker) onto a standard size bottom agar plate and adding 2.5 ml of molten 52°C top agar as in the bait assay, swirling to disperse the host into the top agar and form a lawn, and streaking-in phage with a toothpick either from the plaque archive or directly from well-separated plaques in overlays from the previous step in serial purification. Following plaque formation on the third serial passage plate plaque plugs were picked using barrier tip 1 ml pipettes and ejected into 250 ul of 2216MB to elute overnight at 4°C. Plaque eluates were spiked with 20 ul of host culture and grown with shaking for several hours to generate a primary small-scale lysate. Small scale primary lysates were centrifuged to pellet cells and titered by drop spot assay to estimate optimal inoculum volume to achieve confluent lysis in a 150 mm agar overlay plate lysate. Plate lysates were generated by mixing 250 ul of overnight host culture with primary lysate and plating in 7.5 ml agar overlay. After development of confluent lysis of lawns as compared against negative control without phage addition, the lysates were harvested by addition of 25 ml of 2216MB, shredding of the agar overlay with a dowel, and collection of the broth and top agar. Freshly harvested lysates were stored at 4°C overnight for elution of phage particles, the following day lysates were centrifuged at 5,000 rcf for 20 min and the supernatant filtered through a 0.2 um Sterivex filter into a 50 ml tube and stored at 4°C.

#### Viral genome sequencing

For DNA extraction approximately 18 ml of phage lysate was concentrated using a 30kD centrifugal filtration device (Millipore, Amicon Ultra Centrifugal Filters, Ultracel 30K, UFC903024) and washed with 1:100 2216MB to reduce salt concentrations inhibitory to downstream nuclease treatments. Concentrates were brought to approximately 500 ul using 1:100 diluted 2216MB and then treated with DNase I and RNase A for 65 minutes at 37°C to digest unencapsidated nucleic acids. Nuclease treated concentrates were extracted using an SDS, KOAc, phenol-chloroform extraction and resuspended in EB Buffer (Qiagen, Buffer EB - Elution Buffer, cat# 19086) for storage at 20°C. Phage genomic DNA was sheared by sonication in preparation for genome library preparation. DNA concentrations of extracts were determined using Picogreen (Invitrogen, Quant-iT PicoGreen dsDNA Reagent and Kits, cat# P7589) in a 96-well format and samples brought to 5ug in 100 ul final volume of PCR-grade water diluent for sonication. Samples were sonicated in batches of 6 for 6 cycles of 5 minutes each, at an interval of 30 seconds on/off on the Low Intensity setting of the Biogenode Bioruptor to enrich for a fragment size of ∼300bp. Illumina constructs were prepared from sheared DNA as follows: end repair of sheared DNA, 0.72x/0.21x dSPRI size selection to enrich for ∼300bp sized fragments, ligation of Illumina adapters and unique pairs of forward and reverse barcodes for each sample, SPRI clean up, nick translation, and final SPRI clean up (Rodrigue et al., 2010). Constructs were enriched by PCR using PE primers following qPCR-based normalization of template concentrations. Enrichment PCRs were prepared in octoplicate 25 ul volumes, with the recipe: 1 ul illumina construct template, 5 ul 5x Phusion polymerase buffer, 0.5 ul 10mM dNTPs, 0.25 ul 40uM IGA-PCR-PE-F primer, 0.25 ul 40uM IGA-PCR-PE-R primer, 0.25 ul Phusion polymerase, 17.75 ul PCR-grade water. PCR thermocycling conditions were as follows: initial denaturation at 98°C for 20 sec; batch dependent number of cycles of 98°C for 15 sec, 60°C for 20 see, 72°C for 20 sec; final annealing at 72°C for 5 min; hold at 10°C. For each sample 8 replicate enrichment PCR reactions were pooled and purified by 0.8x SPRI clean up. Each sample was then checked by Bioanalyzer (2100 expert High Sensitivity DNA Assay) to confirm the presence of a unimodal distribution of fragments with a peak between 350-500bp. Sequencing of phage genomes was distributed over 4 paired-end sequencing runs as follows: HiSeq library of 18 samples pooled with 18 external samples, 3 MiSeq libraries each containing ∼100 multiplexed phage genomes. Viral genomes were assembled as previously described^24^. Accession numbers for all sequenced phage genomes are provided in Supplementary Data File 1, where they are identified as “phageSet283”; the subset of phages used in the majority of analyses in this work are identified as “phageSet248” and exclude non-independent isolates derived from the same plaque, as well as well as identical phages isolated from multiple independent plaques from the same host strain in the bait assay.

#### Viral protein clustering

To characterize and annotate groups of proteins in viral genomes in the Nahant Collection proteins were clustered using MMseqs2 v. 2.23394 ^56^ with default parameter settings, the 21,937 proteins reported in the GenBank files associated with each of the 283 Nahant Collection phage genomes were clustered into 5,929 clusters including 2,978 singletons. MMseqs2 cluster assignments for each protein sequence are provided in Supplementary Data File 4.

#### Viral protein cluster annotation

All proteins were annotated using InterProScan^57^ v.5.39-77.0, eggNOG-mapper^58, 59^ v.2 using both automated and viral HMM selection options; with Meta-iPVP^60^; and with best matches to 9518 Viral Orthologous Groups^61^ hmm profiles (obtained at http://dmk-brain.ecn.uiowa.edu/pVOGs/downloads.html); search was performed with hmmer, requiring a bitscore of 50 or greater (highest e-value 5.80E-13), as follows: *hmmsearch -o$out_dir/$hmm_group.$hmmfile.$prots_short_name.hmm.out --tblout$out_dir/$hmm_group.$hmmfile.$prots_short_name.hmm.tbl.out --noali -T 50 $hmmfile$prots_dir/$prots_file*. Annotations for viral protein clusters are provided in Supplementary Data File 4.

______Assignment of proteins to broad functional categories of “Intracellular” acting, “Virion” associated, “Packaging/Assembly” associated, and “Endo/Exolysis” associated was performed at the protein cluster level by manual review of concatenated annotations of all member proteins in the cluster. For select proteins additional characterizations were made using PSI-BLAST^62, 63^, the MPI Bioinformatics implementation of HHpred^64^, and EMBL-EBI implementation of JACKHMMER^65^. All assignments, and concatenated annotations on which they were based, are provided in Supplementary Data File 4.

_____Receptor binding proteins (RBPs) were annotated as follows. RBPs were defined here to include both globular and fibrous host interacting proteins and general protein annotations were reviewed for similarity to known phage receptor binding proteins and supplemented with Phyre2^66^, HHpred, and literature review^67^. Annotated RBPs were mapped onto phage genome diagrams and additional RBPs were annotated based on gene order conservation with phages in the same genus for which RBPs were already identified; annotated RBPs were then used to iteratively search against all Nahant Collection phage proteins using the jackhmmer search tool in the HMMER^68^ v.3.2.1 package (*jackhmmer --cpu 16 -N 3 -E 0.00001 --incE 0.01 --incdomE0.01 -o $run.$1.vs.$2.jackhmmer.iters-$iters.out --tblout $run.$1.vs.$2.jackhmmer.iters-$iters.tbl.out --domtblout $run.$1.vs.$2.jackhmmer.iters-$iters.dom.tbl.out $queryFASTAS$subjectFASTAS*) and new hits were manually reviewed. All annotations were performed on a protein-cluster level and annotations of proteins and protein clusters as “adsorption - RBP” are indicated in Supplementary Data File 4.

_____Recombinases were annotated as follows: Homologs of single strand annealing protein recombinases in the Rad52, Rad51 and Gp2.5 superfamilies in the Nahant Collection phages were identified as follows. First, iterative HMM searches were performed against the Nahant Collection phage proteins using as seeds 194 recombinases identified in Lopes et al.^38^ (excluding RecET fusion protein YP_512292.1; http://biodev.extra.cea.fr/virfam/table.aspx), these represent 6 families of SSAP recombinases (UvsX, Sak4, Sak, RedB, ERF, and Gp2.5); searches were performed using the jackhmmer function of HMMER v.3.1.2 (*jackhmmer --cpu 16-N 5 -E 0.00001 --incE 0.01 --incdomE 0.01 -o $run.$1.vs.$2.jackhmmer.out --tblout$run.$1.vs.$2.jackhmmer.tbl.out --domtblout $run.$1.vs.$2.jackhmmer.dom.tbl.out$queryFASTAS $subjectFASTAS*) – this yielded 156 proteins. Second, all hits were plotted onto genome diagrams for all phages in the collection and additional candidate recombinases identified based on gene neighborhood comparisons – this step identified 4 additional protein clusters (mmseqs 297, 149, 2211, and 600), totaling 224 proteins. Third, all proteins clusters were curated by examination of additional annotations made using InterProScan^57^, EggNOG-mapper^58^, Phyre2^66^, and HHpred ^64^. Following this third step, there were 3 protein clusters for which support was limited, these were included in the final dataset as putative SSAP recombinases but are highlighted here. **Protein cluster mmseq 297** (present in 21 phages in 6 genera): was always encoded by genes adjacent to genes in protein cluster mmseq 3923, which was itself a recombinase associated exonuclease that was found either adjacent to mmseq 297 or to the well-supported putative SSAP recombinase mmseq 3721 (sometimes separated by one gene from mmseq 3721). **Protein cluster mmseq 600** (present in 2 phages in 2 genera): was encoded adjacent to a protein cluster annotated as a recombination associated exonuclease; iterative HHMER searches of a mmseq 600 cluster representative (AUR82881.1) against Viruses in UniProtKB using jackhmmer yielded hits to proteins in mmseq 297 in iteration 3. **Protein cluster mmseq 2990** (present in 1 phage): was encoded adjacent to two small proteins encoding putative recombination associated exonucleases and was in the same genomic position relative to neighboring genes as putative recombinases in related phages in the genus. Finally, all putative SSAP recombinase genes were assigned to a recombinase family by clustering based on 2 iterations of all-by-all HMM jackhmmer sequence similarity searches of all candidates and the reference seed set of Lopes^38^ (*jackhmmer --cpu 16 -N 2 -E 0.00001 --incE 0.01 --incdomE 0.01 -o $run.$1.vs.$2.jackhmmer.out --tblout$run.$1.vs.$2.jackhmmer.tbl.out --domtblout $run.$1.vs.$2.jackhmmer.dom.tbl.out$queryFASTAS $subjectFASTAS*); similarities were were visualized using CytoScape v.3.3.0 using the “Edge-weighted Spring Embedded Layout” based on jackhmmer score (Supplementary Figure [recombinases]); clusters were identified using the ClusterMaker2 v.1.2.1 Cytoscape plugin with the MCL cluster option and all settings at default and Granularity=2.5. Proteins in 3 mmseq clusters did not fall into MCL clusters with recombinases from the annotated seed set and therefore are described as “unknown” rather than being assigned to a family of recombinases. All final assignments of genes to a recombinase superfamily and family, as well as all associated annotations, are provided in Supplementary Data File 4.

___Annotation of viral potential for temperate lifestyle was conducted as follows.To examine whether any of the phages in the collection are potentially temperate we searched for characteristic lysogeny associated genes and performed BLAST searches against the 294 bacterial genomes included in the host range assay. The sole phage in NCVG_31 is a prophage directly derived from its host of isolation, the only such case. The phages in NCVG_41 are identified as Mu-like phages, with similar prophages present in the 294 bacterial genomes screened. The phages in NCVG_12 (2 phages), NCVG_24 (4 phages), and NCVG_31 (1 phage), were identified as encoding integrases (based on annotations with iterative Jackhmmer searches with PF00589 phage integrase seed alignment and with EggNOG-mapper). Iterative Jackhmmer searches with N15 phage linear plasmid maintenance gene SopA (NP_046923.1) and *E. coli* ParA (AAA99230.1) yielded no hits.

#### Viral genome clustering

To understand how the diversity of viral genomes in the Nahant Collection is organized with respect to units equivalent to those representative of genera previously defined by the International Committee on the Taxonomy of Viruses (ICTV) we use the VICTOR classifier^32^, which determines genome to genome distances between concatenated amino acid sequences of viral proteomes using the Genome-BLAST Distance Phylogeny method^69^ and clusters these using OPTSIL^70^ and criteria optimized by benchmarking to ICTV prokaryotic virus taxonomic units^32^ with the fraction of links required for cluster fusion of 0.5^71^. Average support values for the phylogenomic trees using the D0, D4, and D6 VICTOR formulas were 49%, 31%, and 51%, respectively, and results presented here were those derived from the D6 formula, for which 171 species-level and 49 genus-level clusters. VICTOR taxonomy assignments for all phages included in the analysis are provided in Supplementary Data File 1B, where they are identified as “phageSet283”.

#### Viral genome relatedness to previously described phages

To assess the robustness of the predicted VICTOR-based genus level groupings and their relation to previously identified viruses with known hosts we clustered the Nahant Collection phages with >10,000 previously described phage genomes. We use the curated list of NCBI phage genomes generously provided as a public resource by the laboratory of Andrew Millard (16103 phage genomes; http://s3.climb.ac.uk/ADM_share/crap/website/26Aug2019_phages.gb.gz). We reduced the full list of 16,103 phage genomes to 10,663 genomes based on 95% identity clustering using the dedupe.sh tool in BBtools; we then added Nahant Collection phages (identified as “phageSet283” in Supplementary Data File 1B) not already included in the list, yielding a total of 10,722 phage genomes (see Supplementary Data File 2A for a list of all phage genome accessions included). We next used vConTACT2 v.0.9.10^29^ to predict viral genome clusters (*vcontact --raw-proteins $rawprots --rel-mode ’Diamond’ --proteins-fp $gene2genome --db ’ProkaryoticViralRefSeq94-Merged’ --pcs-mode MCL --vcs-mode ClusterONE --c1-bin/home/k6logc/miniconda3/bin/cluster_one-1.0.jar --output-dir $outdir*; see Supplementary Data File 2B for full clustering output for all members). Correspondence between VICTOR genera and vConTACT2 clusters was overall high (see Supplementary Data File 2C for cluster assignments for Nahant phages and Supplementary Data File 2D for correspondence between genera and vConTACT2 clusters). All vConTACT2 subclusters included phages from only 1 VICTOR genus, however, 3 VICTOR genera were split across vConTACT2 clusters or subclusters. The cases of VICTOR genera split by vConTACT2 included: NCVG_17 (the *Autolykiviridae*), split into 2 vConTACT2 clusters; NCVG_36, split into 2 vConTACT2 clusters; NCVG_23, split into 2 vConTACT2 outliers; and NCVG_47, with all members within one vConTACT2 cluster but split across 3 subclusters. Three VICTOR genera (NCVG: 14, 21, and 46) that otherwise corresponded with vConTACT2 clusters each included a member classified as an outlier by vConTACT2. And, 7 phages (members of NCVG: 7, 9, 12, and 28) that were included as inputs to the vConTACT2 analysis were excluded from the summary outputs in multiple run attempts for reasons that are unclear. The vConTACT2 analysis allowed us to identify 47 phages previously described phages that belong to 17 Nahant Collection genera; none of these previously described phages were identified as members of the same species as any Nahant phages when we re-evaluated these 17 genera with the VICTOR classifier with all members included (Supplementary Data File 2E for information about 47 previously described phages). As noted in the main text, the majority of previously described phages in VICTOR genera with Nahant Collection phages also infected hosts in either the Vibrionaceae or Shewanellaceae (a second family of hosts also represented in this study), consistent with previous finding that phage genera are specific to host families^32^. However, in 4 of the 17 genera, previously described phages had hosts in non-Vibrionales Gammaproteobacterial orders, including the Enterobacterales, Aeromonadales, Pasteurellales, and Alteromonadales. The genus of phages for which previous isolates had the most diverse hosts (NCVG_31) contains phages that, in the 2019 taxonomy revision by the International Committee on the Taxonomy of Viruses were assigned to multiple genera within the phage subfamily Peduovirinae; and was represented in the Nahant Collection by only a single phage - the sole case in this study of isolating a host-derived prophage (as the result of a prophage forming a plaque on its own host in the bait assay).

### Host range

#### Host range assay

Host range of viruses was determined as follows, and as also previously described^22, 28^. Cell-free phage lysates were stamped onto host agar overlay lawns and observed for changes in lawn morphology proximal to each stamp. Phage application to host lawns was performed using a 96-well blotter (BelArt, Bel-blotter 96-tip replicator, cat # 378760002) that was set into a microtiter plate containing arrayed phage lysate, transferred to the surface of the host lawn, and allowed to remain in contact for several minutes. Each 96-stamp contained 3 replicates of each phage lysate, distributed across three panels (columns 1-4, 5-8, 9-12) each with a unique array of the 32 samples (including one negative control). 96-well blotters were microwave steam sterilized (Tommee Tippee, Closer to Nature Microwave Steam Sterilizer) in batches for continuing re-use during plating sessions. Bacterial strains were prepared for the infection assay by inoculating 1ml volumes of 2216MB in 2ml 96-well culture blocks directly from glycerol stocks and shaking them at RT for approximately 48 hours. Agar overlays were prepared by transferring 250 ul aliquots of host culture to bottom agar plates (2216MB, 1% Bacto Agar, 5% glycerol; in 150mm diameter plates) and adding 7ml of molten 52°C top agar (2216MB, 0.4% Bacto Agar, 5% glycerol). Phages were prepared by distributing lysates into a 2ml 96-well culture block in panels as described above, aliquots of <200 ul were then transferred into shallow microtiter plates so that the blotter could phage lysate by capillary action. Host lawns were stamped with phage lysates within 5-6 hours of plating lawns. Agar overlays were assessed for changes in lawn morphology associated with phage treatment and scored blind with respect to phage identity and arrangement of replicates. Plates were scored for the presence of interactions on days 1, 2, 3, 7, 14, 21, and 30, and the outer bands of the interaction zones were marked with a different color for each time point. After 30 days the interactions for each strain were recorded and the approximate diameter for each interaction at each time point was recorded. During recording of the interactions for each plate an additional qualitative measure of confidence in the projected positive or negative call of the interaction was made. For example, where 2 of 3 replicates were positive for a phage on a lawn with no other positive interactions such an interaction would be called by the qualitative measure as “real”; alternatively, where 2 of 3 replicates were positive for a phage on a lawn containing several other positive interactions the qualitative measure might call these replicates “contam” if they were high-titer interactions and occurred in close proximity to other positive interactions. List of all infection pairs are provided in Supplementary Data File 1, and represent interactions between phages identified in the Supplement as “phageSet248” and hosts identified as “baxSet279”.

### Characterization of phage-host interaction features

#### BiMat modularity analysis

To characterize large scale features of the infection network we use the BiMat MatLab package^27^ and as described in^2,15, 72^. Modularity was quantified using the leading eigenvector method, with a Kernighan-Lin tuning step performed after module detection, and nestedness was quantified using overlap and decreasing fill (NODF). Statistical significance of modularity and nestedness was tested against 1000 random matrices generated using the equiprobable method, preserving the overall matrix connectivity. Modularity values were as follows: Qb value 0.7306, mean 0.4362, std 0.0047, z-score 63.2774, t-score 2001.0077, percentile 100; Qr value 0.9318, mean 0.1004, std 0.0219, z-score 37.9184, t-score 1199.0848, percentile 100. Nestedness values were as follows: Nestedness value: 0.0300, mean 0.0230, std 0.0005, z-score 14.0305, t-score 443.6833, percentile 100. The 248 phages included in the analysis (“phageSet248 in the Supplementary Data) are genome sequenced phages isolated during the Nahant study, excluding cases of duplicate phages purified from the same plaque and excluding duplicate phages purified from different plaques on the same host; the 279 bacteria included in the analyses (“baxSet279” in Supplementary Data File 1) include all bacterial strains screened in the host range assay for which there was a positive interaction with a phage in phageSet248 (ie. host strains that were assayed but not killed by any phages were not included); these represent 1,436 infections out of a possible set of 69,192 and yield a connectance or fill of 0.021. To facilitate visual comparisons between the matrix of interactions between phages and bacteria with known species assignments (Fig. 2b) and the BiMat results, the representation of the BiMat analysis as shown in the main text (Fig.2a) includes only the subset of interactions between phages in phageSet248 and the 259 bacterial strains (“baxSet259”) that is the intersection of the 279 bacterial strains (“baxSet279”) included in the BiMat analysis and the 294 bacterial genomes (“baxSet294”) for which genomes were available. BiMat module assignments for all phages and hosts are provided in Supplementary Data File 1 sheet C.

#### Average nucleotide identity

FastANI v.1.3 was used to determine average nucleotide identity (ANI) for phages and hosts as follows. For phages, run parameters were: kmer size of 16, fragment length of 100, minimum fraction of shorter genome coverage of 75% (*fastANI -t 12 -k 16 --fragLen 100 --minFraction 0.95 --matrix --ql $genomesPathsList --rl $genomesPathsList -o phageSet248_k16fL100Frax95.fastani.out*). For bacteria, run parameters were: kmer size of 16, fragment length of 3000, minimum fraction of shorter genome coverage of 50% (*fastANI -t 12 -k 16 --fragLen 3000 --minFraction 0.5 --matrix --ql $genomesPathsList --rl $genomesPathsList -o baxSet294_k16fL3000Frax50.fastani.out*). Results of ANI analyses are provided in Supplementary Data File 1.

#### Host range divergence analyses within species and genera

To quantify overlap in host range profiles between phages we develop a metric of host range divergence (represented as concordance (1-divergence) in the main text and Fig. 3). We normalize the binary vector x_i_ = (*x*_1_, *x*_2_, …, *x_m_*) representing the killing host range of a phage *i* across all *m* hosts so that it sums to 1, and interpret the result 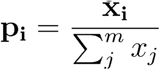 as a probability distribution of killing across all *m* hosts for a single phage *i*. We then define the *scaled host range divergence* of a given genus consisting of *n* phages to be the normalized generalized Jensen-Shannon divergence (gJSD) of their infection probability vectors, 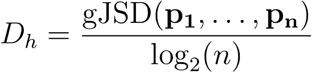. This has the property that 0 ≤ *D_h_* ≤ 1, where *D_h_* = 0 means the host ranges of all phages overlap exactly, and *D_h_* = 1 means none of the host ranges have any overlap with each other. We note that we took a conservative approach in determining genus-level concordances as presented in Fig. 3 by including all phages within each genus in the calculation of concordance rather than collapsing intra-species blooms, which are largely comprised of phages with overlapping host ranges. Because high overlap of host range within species increases the value of the overall genus-level concordance metric (because of the higher number of pairwise comparisons with high values of concordance within species), this approach will be affected by the evenness of species abundances within a genus and including all members yields up to >24 times higher values of the concordance metric than when only including single representatives of each species in this dataset. Calculations of concordance are provided in Source Data Figure 3 for genera using both the approach of using all members (Source Data Figure 3 sheet B, with phages identified as “phageSet248”) and that of using only single representatives from each species (Source Data Figure 3 sheet C, with phages identified as “phageSet171”); values for species are provided in Source Data Figure 3 sheet A.

### Characterization of sequence sharing

#### Homologous recombination within species and genera

To assess the extent of recombination between closely related viruses we used viral species and genera as operationally defined by VICTOR as the framework and estimated effective relative contribution of recombination over mutation (r/m) as follows. HomBlocks v.1.0 ^73^ was used with default parameter settings to identify, extract, and trim conserved regions within genera based on alignments with progressiveMauve (build February 13, 2015)^74^ and trimAl v.1.2^75^. Phylogenetic relationships between sequences were predicted using IQ-TREE v.1.6.12^76^. ClonalFrameML v.1.12^77^ was then used to evaluate evidence for recombination within species and genera. Estimates of relative effect of recombination to mutation (r/m) were based on the formula r/m = R/theta * delta * nu; where R/theta is the ratio of recombination to mutation rate, delta is mean import length, and nu is the nucleotide distance between imported sequences. We include in our analysis a control set of closely related siphovirus genomes which in a previous study were found to have an r/m of 23.50^34^, accession numbers for these phages are provided in Source Data Figure 3 sheet G; using the methods we describe here we find a similar r/m of 18.3. All analyses were performed only where there were ≥3 phages per species or genus. For genus level calculations of r/m we performed analyses using two different sets of genomes: first, as presented in Fig. 3e,g including all phages in each genus (Source Data Figure 3 sheet E, using phages identified as “phageSet248”); second, including only 1 representative from each species (Source Data Figure 3 sheet F, using phages identified as “phageSet171”). As for estimates of host range divergence, estimates of r/m are affected by evenness of species abundances within a genus and including all members can result in reduced estimates of r/m; for example, for NCVG_17, the *Autolykiviridae*, estimates of genus-level r/m are >285 higher when considering only species representatives rather than all members of the genus.

#### Sharing of 25-mers

To assess nucleotide sequence sharing between phages in the collection overall, we used methods that were not limited by requirement for large scale pairwise genome conservation. We use Mash v.2.2.2^78^ to create a sketch file for all phage genomes (*mash sketch-o $out -k 400000 -s 25 -i $concatGenomesFile*), using a k-mer size of 25 and a sufficiently large sketch size to capture all 25-mers in the largest phage genome; we next generate the mash distance output (*mash dist $out.msh $out.msh -i >> $out.msh.ALLbyALL.dist*), which includes a count of shared 25-mers between all pairs of phage genomes. To compare the infection and k-mer sharing networks, we created a binary vector reflecting whether each pair of phages shares at least one host in the observed infection matrix. Similarly, we created a binary vector reflecting whether each pair of phages shares at least one 25-mer between their respective genomes, as reported by Mash. We then calculated the mutual information between these vectors using the R function mutinformation() from package infotheo. Using mutual information analysis to ask whether phages that share ≥1 25-mer also share ≥1 hosts, we found that one matrix does not predict the other (0.018 bits out of a maximum value of 1 predicted).

Host sharing information and Mash outputs for all phage-phage pairs are provided in Source_Data_Figure_4 and were based on analysis of phages identified in Supplementary Data File 1 as “phageSet248”.

#### BRIG plotting

To visualize regions of genome conservation and divergence within sets of phages as compared to a reference, as shown in Fig. 4d we used the BLAST Ring Image Generator tool BRIG^79^.

#### Transfers with identifiable directionality between genera

To use a method that could predict directionality of transfer between phages in different genera, we used MetaCHIP^80^, providing VICTOR genus as the grouping variable. This method is a conservative estimator as it requires that candidate regions be conserved within donor groups but not in recipient groups. Modifications were performed to the source code to accept the amino acid-based phylogenetic tree as output by VICTOR as the phylogeny used in the MetaCHIP analysis. MetaCHIP parameters were set to 90% identity cutoff, 200bp alignment length cutoff, 75% coverage cutoff, 80% end match identity cutoff, and 10Kbp flanking region length. MetaCHIP results are provided in Source_Data_Figure_4.

#### Potential for cross-recruitment of reads between viral genomes

To assess the potential for cross-recruitment of metagenomic reads to lead to false positive detection of phage genomes in metagenomic sequencing datasets we simulated Illumina reads for each of the phage genomes using parameters representative of high quality viral metagenome datasets and then asked whether they yielded a positive identification using a recently developed reference mapping tool designed for rapid characterization of large phage metagenomes. We included 248 phages of the Nahant Collection, as well as 47 previously described phage genomes that in vConTACT2 analyses (above) were found to be members of the same VICTOR D6aa genera as the Nahant Collection phages. Simulated Illumina paired end reads were generated using DWGSIM v.1.1.11^81^, with settings of: outer distance 500, standard deviation of read lengths 50, number of reads 100000, read lengths for both reads of 250bp, and error rate 0.0010 (*dwgsim -d 500 -s 50-N 100000 -1 250 -2 250 -r 0.0010 -c 0 -S 2 $genome $genome.simreads*). Cross recruitment of reads to each of the 295 phages was performed independently for simulated reads from each phage using default settings of FastViromeExplorer^82^ for criteria required to define a genome as present in a read dataset: genome coverage [0.1]; ratio of coverage to expected coverage [0.3], which considers evenness of read mapping; and minimum number of mapped reads [10]. Notably, performing this analysis with 100bp-length read sets eliminated cross-genus mappings that were identified using 250bp read sets. This is thought to occur because FastViromeExplorer uses a Kallisto^83^ based pseudoalignment approach, which requires only a single matching 31-mer to map a given read to a target genomes, and thus where a 100bp read and a 250bp read both share only a single 31-mer with a reference genome the 250bp read will result in overall greater coverage of the genome. All information about the phages included in the cross-recruitment analysis, and the associated data, are provided in Source_Data_Extended_Data_Figure_5

